# Repetitive magnetic stimulation induces plasticity of excitatory synapses through cooperative pre- and postsynaptic activity

**DOI:** 10.1101/2024.11.04.621890

**Authors:** Christos Galanis, Maximilian Lenz, Mohammadreza Vasheghani Farahani, Andreas Vlachos

## Abstract

Transcranial magnetic stimulation (TMS) is a widely used non-invasive technique in research and clinical settings. Despite its success, the cellular and molecular mechanisms underlying TMS-induced changes in the brain remain incompletely understood. Current protocols are largely heuristic, based on system-level observations. This study employed in vitro repetitive magnetic stimulation (rMS) in mouse brain tissue cultures, combined with computational modeling to develop an experimentally validated approach for predicting TMS effects. Unlike electrical or optogenetic stimulation, rTMS uniquely enhances plasticity by activating both pre- and postsynaptic neurons, with brain-derived neurotrophic factor (BDNF) playing a crucial role. Our simulations accurately predicted the frequency-dependent effects of rTMS, providing a critical step towards developing robust, validated tools that will enhance the precision and effectiveness of TMS applications across both research and clinical settings.

**Significance Statement:** This study elucidates the mechanisms by which TMS promotes neural plasticity, specifically long-term potentiation (LTP) of excitatory neurotransmission. By demonstrating that rMS engages both pre- and postsynaptic neurons with BDNF as a key mediator, it provides a mechanistic basis for the therapeutic effects of rTMS. These insights advance our understanding of rTMS-induced brain plasticity and support the development of predictive computer models, paving the way for more effective and standardized TMS protocols in research and clinical applications.

## Introduction

Transcranial magnetic stimulation (TMS) has been extensively studied for its ability to modulate excitability of the human cortex beyond the stimulation period (1–3). Based on the physical principle of electromagnetic induction, TMS allows for the non-invasive stimulation of cortical networks through the intact skin, skull, and meninges. Meanwhile, it is well established that repetitive TMS (rTMS), i.e., application of several hundred TMS-pulses, induces long-lasting changes in synaptic transmission (4, 5). Yet, the precise cellular and molecular mechanisms underlying rTMS-induced plasticity are not well-understood. As a consequence, it is challenging to predict the outcome of a given stimulation protocol, which poses significant challenges in controlling and standardizing the “biological dose” of rTMS across brain regions and subjects (6, 7).

Studies in animal models or in appropriate *in vitro* preparations–combined with computational modeling–provide suitable experimental approaches in this context, since basic science techniques can be used to unravel how repetitive magnetic stimulation (rMS) induces neural plasticity. In our previous work we were able to demonstrate that rMS of mouse and rat brain tissue cultures induces long-lasting structural and functional changes of excitatory postsynapses (4, 8, 9). A 10 Hz rMS protocol consisting of 900 pulses induced robust potentiation of excitatory inputs on CA1 pyramidal neurons (8, 9). Since these changes were mediated by N-methyl-D-aspartate receptors (NMDARs), the results of these earlier studies suggested that 10 Hz rMS is capable of inducing NMDA receptor dependent long-term potentiation (LTP) of α-amino-3-hydroxy-5-methyl-4-isoxazolepropionic acid receptor (AMPAR)-mediated synaptic transmission (4, 9). The induction of LTP using a 10 Hz stimulation protocol applied through local electrical stimulation is not anticipated (10, 11). Coordinated pre- and postsynaptic activation, often through paired stimulation, is essential for achieving LTP at lower frequencies (12, 13).

In the present study, we compared the effects of 10 Hz rMS with local electrical and optogenetic stimulation employing the exact same 10 Hz stimulation protocol. While we confirmed previous findings on rMS-induced potentiation of excitatory neurotransmission, both local electrical and optogenetic stimulation of afferent inputs only resulted in long-term depression (LTD). Experiments utilizing chemogenetic tools and Ca^2+^ imaging during stimulation demonstrated that rMS facilitates plasticity through the activation of both pre- and postsynaptic neurons. Computational modeling based on spike-timing dependent plasticity (STDP) effectively replicated the outcomes of 10 Hz rMS and predicted the frequency-dependency of our 900 pulse protocol in the experimental context. Furthermore, we demonstrated that BDNF is crucial for 10 Hz rMS-STDP: in the presence of a soluble scavenging receptor, 10 Hz rMS could not trigger LTP, while pharmacological activation of tyrosine receptor kinase B (TrkB)/BDNF signaling converted 10 Hz electrically induced LTD into LTP. These results advance our understanding of how rTMS achieves lasting changes in neural transmission and cortical excitability. They indicate that BDNF release, stimulated by rTMS-STDP in targeted brain regions, may contribute to the beneficial effects of rTMS observed in clinical settings.

## Results

### 10 Hz repetitive magnetic stimulation (rMS) potentiates excitatory synapses

Tissue cultures (≥18 days *in vitro*) including the entorhinal cortex and the hippocampus were stimulated with a 10 Hz stimulation protocol comprising 900 pulses (divided into 9 sets of 100 pulses with 30-second inter-train-intervals). The stimulation was delivered with a standard 70 mm figure-of-eight coil (Figure 1*A, B*). In line with previous findings (8, 9), whole-cell patch-clamp recordings from CA1 pyramidal neurons (Figure 1*C*) demonstrated a significant increase in mean amplitude of AMPA receptor-mediated miniature excitatory postsynaptic currents (mEPSCs) 2 – 4 h after 10 Hz stimulation (Figure 1D, E). These results are consistent with a potentiation of glutamatergic synapses.

**Figure 1.**
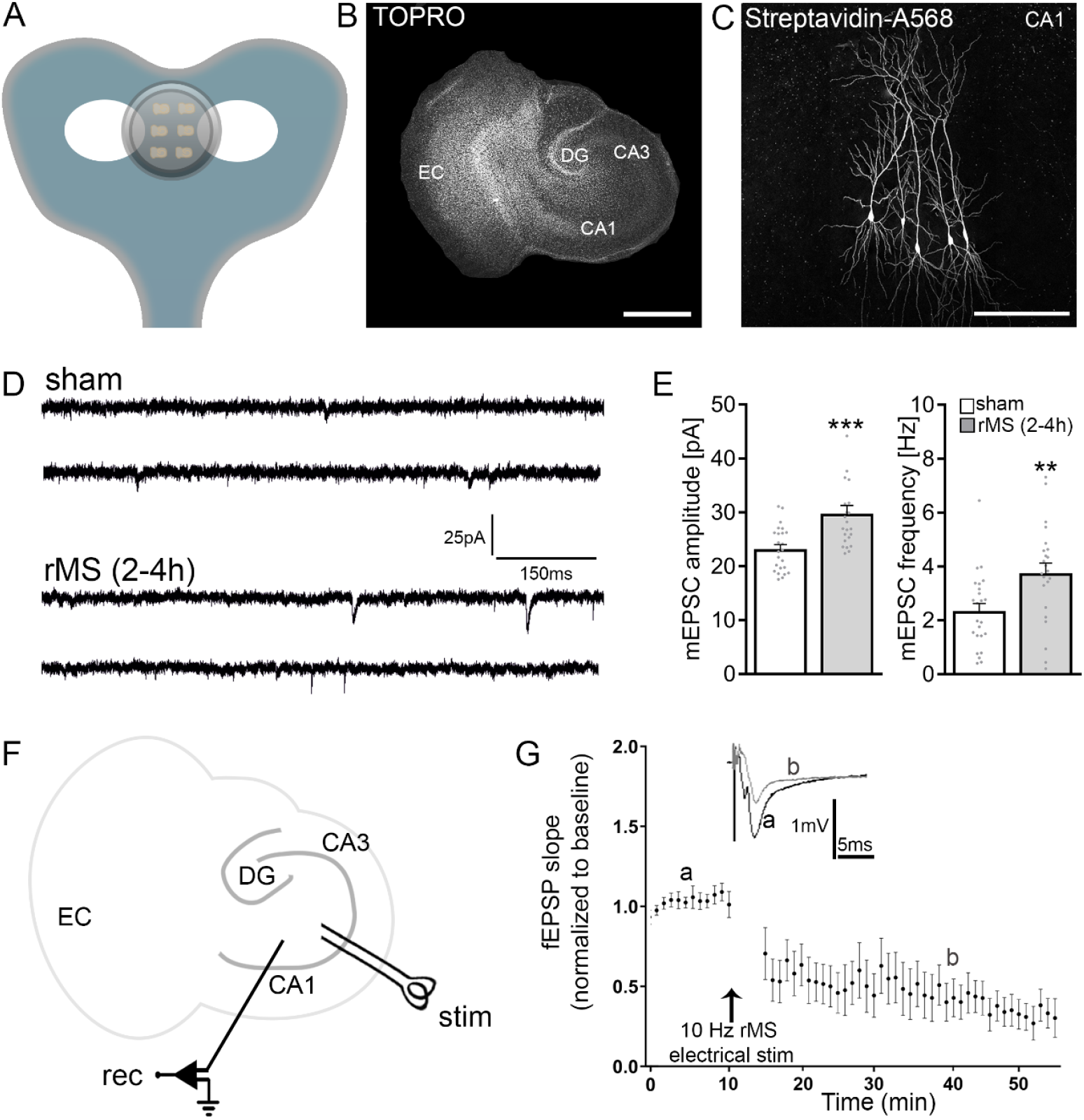
10 Hz repetitive magnetic stimulation (rMS) induces synaptic plasticity in mouse organotypic tissue cultures (A) Schematic illustration of the experimental setting. Organotypic tissue cultures are stimulated in a standard 35 mm petri dish filled with standard extracellular solution with AirFilm® Coil (900 pulses, 10 Hz, at 50 % maximum stimulator output). (B) Overview of an organotypic tissue culture. Visualization of cytoarchitecture with TOPRO. DG, Dentate gyrus; EC, entorhinal cortex; CA1 and CA3, *Cornu Ammonis* areas 1 and 3. Scale bar, 500□μm. (C) Patched CA1 pyramidal neurons filled with biocytin and identified *post hoc* with streptavidin-A568. Scale bar, 100□μm. (D, E) Sample traces and group data of AMPA receptor-mediated miniature excitatory post synaptic currents (mEPSCs) recorded from mouse CA1 pyramidal neurons in control (sham) and stimulated (rMS) cultures 2 – 4 h post stimulation (n_sham_ = 24 neurons, n_rMS(2-4h)_ = 23 neurons; Mann-Whitney test). (F) Schematic of a classic Schaffer-collateral LTP experiment in organotypic tissue cultures. Stim, stimulation electrode; rec, recording electrode G) Excitatory post synaptic field potential (fEPSP) slopes recorded in the CA1 stratum radiatum following a 10 Hz, 900 pulses electrical stimulation of the Schaffer collaterals (n□=□6 tissue cultures). Individual data points are indicated in this and the following figures by gray dots. Data are mean ± SEM. NS, Not significant. **p□<□0.01, ***p□<□0.001.

### 10 Hz electrical stimulation of Schaffer collateral-CA1 synapses induces long-term depression of excitatory postsynaptic potentials

To further explore the impact of 10 Hz stimulation on synaptic plasticity, we conducted experiments using local electrical stimulation in tissue cultures. A stimulating electrode was placed to activate Schaffer collaterals, facilitating the recording of field excitatory postsynaptic potentials (fEPSP) from CA1 stratum radiatum (Figure 1F). Before applying the identical 10 Hz stimulation protocol comprising 900 pulses (divided into 9 sets of 100 pulses with 30-second inter-train-intervals) baseline recordings were obtained at a frequency of 0.33 Hz for 10 min and changes in fEPSP slopes were assessed. In these experiments, we observed a reduction in fEPSP slopes following 10 Hz stimulation (Figure 1*G*), indicating a consistent long-term depression (LTD) of glutamatergic neurotransmission. We concluded that the same 10 Hz protocol, which induces synaptic potentiation when applied via rMS, leads to LTD when locally applied via electrical stimulation of afferent input.

### Optogenetic stimulation of both CA3 and CA1, or CA3 only, reproduces results observed with rMS and electrical stimulation

We introduced AAV1-hSyn-CrimsonR-tdTomato in CA3, CA1, or both regions through local injections (Figure 2*A*) to explore the impact of non-invasively stimulating pre-, post-, and both synaptic sites simultaneously with light. Neurons were activated using a 525+ nm light source, employing the 10 Hz stimulation protocol. Whole-cell patch-clamp recordings reliably detected light-induced action potentials in CrimsonR-tdTomato-expressing CA1 neurons comprising 900 action potentials divided into 9 sets of 100 pulses with 30-second inter-train-intervals (Figure 2B).

**Figure 2.**
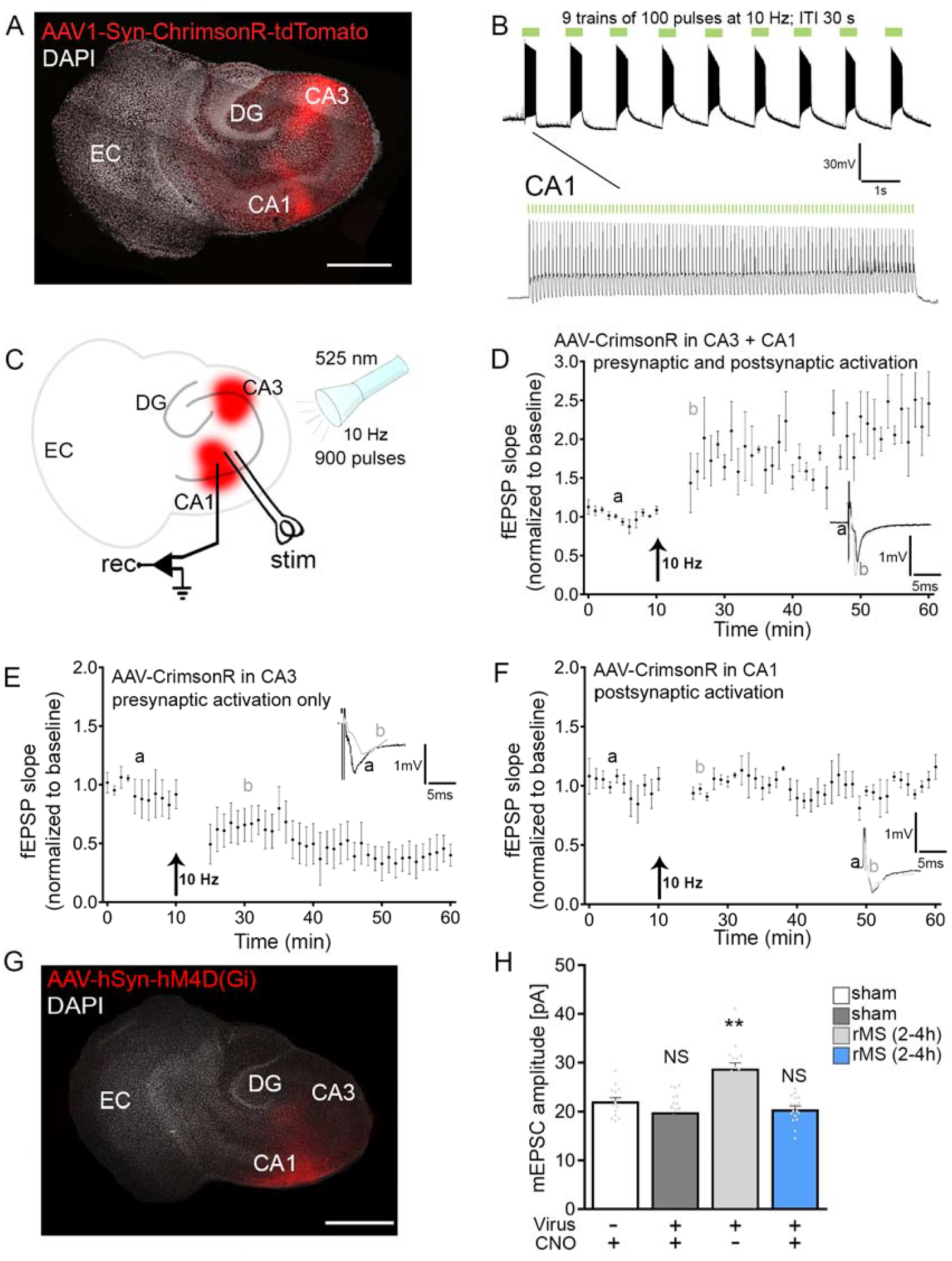
Pre- and postsynaptic activation of excitatory synapses leads to Schaffer collateral LTP and potentiation of CA1 excitatory synapses. (A) Overview of a tissue culture transduced with AAV1-hSyn-CrimsonR-tdTomato in the CA1 and CA3 region. DG, Dentate gyrus; EC, entorhinal cortex; CA1 and CA3, *Cornu Ammonis* areas 1 and 3. Scale bar, 500□μm. (B) Sample trace of light induced responses of a patched CA1 neuron expressing the opsin CrimsonR. (C, D) Excitatory post synaptic field potential (fEPSP) slopes recorded in the CA1 stratum radiatum following a 10 Hz, 900 pulses optical stimulation of the transduced CA3 and CA1 pyramidal neurons (n = 4 tissue cultures) (E) Excitatory post synaptic field potential (fEPSP) slopes recorded in the CA1 stratum radiatum following a 10 Hz, 900 pulses optical stimulation of the transduced CA3 pyramidal neurons (n = 5 tissue cultures). (F) Excitatory post synaptic field potential (fEPSP) slopes recorded in the CA1 stratum radiatum following a 10 Hz, 900 pulses optical stimulation of the transduced CA1 pyramidal neurons (n = 3 tissue cultures). (G) Overview of a tissue culture transduced with AAV1-hSyn-hM4D(Gi)-tdTomato in the CA1 region. DG, Dentate gyrus; EC, entorhinal cortex; CA1 and CA3, *Cornu Ammonis* areas 1 and 3. Scale bar, 500□μm. (H) Group data of AMPA receptor-mediated mEPSCs recorded 2 – 4 h after stimulation from virus expressing, or control, CA1 pyramidal neurons in presence or absence of CNO (n_sham+CNO_ = 15 neurons, n_sham+virus+CNO_ = 20 neurons, n_rms(2-4h)+virus_ = 16 neurons, n_rms(2-4h)+virus+CNO_ = 17 neurons; Kruskal-Wallis test). Data are mean ± SEM. NS, Not significant. **p□<□0.01

Baseline electrically evoked fEPSPs were taken prior to the application of optogenetic 10 Hz stimulation as described above (c.f., Figure 1G), with subsequent changes in fEPSP slopes assessed. In cultures expressing CrimsonR-tdTomato in both CA3 and CA1, the optogenetic 10 Hz stimulation led to synaptic potentiation, indicated by a significant enhancement in the slopes of electrically evoked fEPSPs (Figure 2*C, D*). Conversely, employing the same protocol in slice cultures expressing CrimsonR-tdTomato only in CA3 induced LTD of fEPSP slope (Figure 2*E*), similar to what we observed with local electrical stimulation of Schaffer collaterals (c.f., Figure 1*G*). Optogenetic activation of CA1 alone did not produce significant changes in excitatory neurotransmission (Figure 2*F*). The results prompted the hypothesis that plasticity induced by 10 Hz rMS requires simultaneous activation of both pre- and postsynaptic neurons.

### Chemogenetic silencing of CA1 during 10 Hz rMS does not result in potentiation of glutamatergic neurotransmission

To investigate the importance of coordinated pre- and postsynaptic neuron activation for plasticity induced by 10 Hz rMS, we used a chemogenetic approach to hyperpolarize CA1 neurons during rMS. This involved the local injection of AAV9-hSyn-hM4D(Gi) into CA1 (Figure 2G). The administration of the specific agonist Clozapine-N-Oxide (CNO, 100µM) led to a significant hyperpolarization of the membrane potential, reducing the probability of action potential generation during whole-cell recordings upon 300 pA current injections (data not shown).

We stimulated tissue cultures with 10 Hz rMS in the presence of 100 µM CNO. The previously described rMS-induced increase excitatory synaptic strength was not observed 2 – 4 h post-stimulation (Figure 2H). Cultures transfected with the virus and stimulated with 10 Hz rMS (in presence of vehicle-only) reliably replicate the observed increase in mEPSC amplitude 2 – 4 h after stimulation (Figure 2H). These findings supported the hypothesis that the synergistic activation of pre- and postsynaptic neurons is crucial for the potentiation of excitatory synapses induced by 10 Hz rMS.

### Calcium imaging during 10 Hz rMS shows simultaneous activation of CA3 and CA1 neurons

To further support the pre- and postsynaptic activation by 10 Hz rMS, we conducted Ca2+-imaging during 10 Hz rMS sessions. Whole tissue cultures were transduced with AAV1-hSyn-GCaMP6f and imaged using a microscope fitted with a carbon stage and a standard 70 mm figure-of-eight coil (Figure 3). 10 Hz rMS triggered a synchronized rise in GCaMP6f fluorescent intensity (ΔF/F_0_) of both CA3 and CA1 regions, a phenomenon not observed under baseline network activity (Figure 3A, B). Cultures transduced with GCaMP6f and locally injected with AAV9-hSyn-hM4D(Gi) in CA1, when imaged during 10 Hz rMS, exhibited no significant calcium transients in CA1 in the presence of 100 µM CNO. We concluded that rMS-induced plasticity depends on synchronized depolarization and coordinated activation of pre- and postsynaptic neurons.

**Figure 3.**
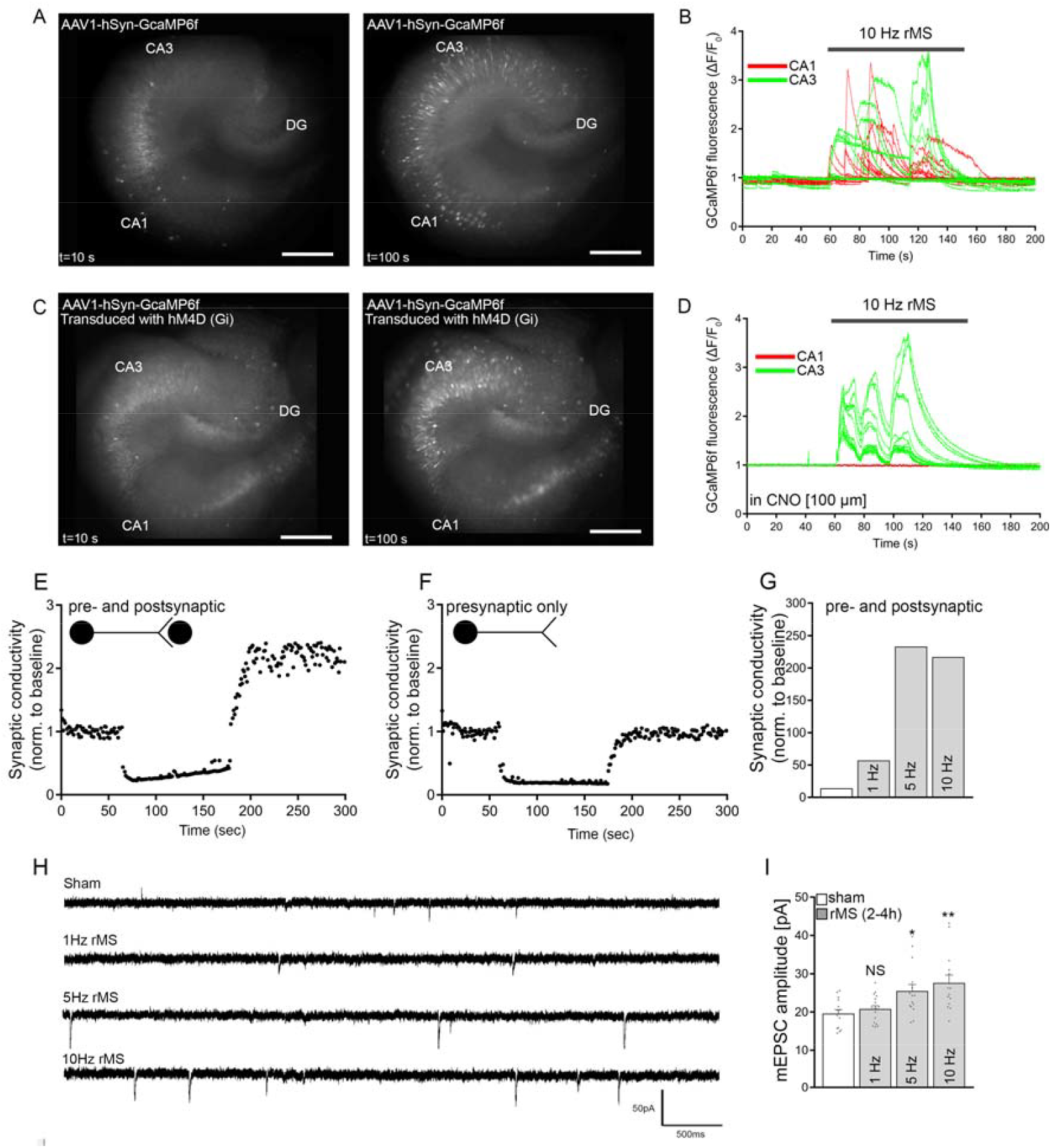
Calcium imaging during rMS indicates simultaneous activation of CA1 and CA3. (A) Overview of an AAV1-hSyn-GCaMP6f transduced tissue culture. DG, Dentate gyrus; CA1 and CA3, *Cornu Ammonis* areas 1 and 3. (B) GCaMP6f fluorescence (ΔF/F_0_) during baseline and rMS (10 Hz, 900 pulses) from 10 CA1 and 10 CA3 pyramidal neurons (n = 3 tissue cultures). (C) Overview of an AAV1-hSyn-GCaMP6f and AAV1-hSyn-hM4D(Gi)-tdTomato transduced tissue culture. DG, Dentate gyrus; CA1 and CA3, *Cornu Ammonis* areas 1 and 3 (n = 2 tissue cultures). (D) GCaMP6f fluorescence (ΔF/F_0_) during baseline and rMS (10 Hz, 900 pulses) from 10 CA1 and 10 CA3 pyramidal neurons in the presence of CNO. (E, F) Computational modeling revealing LTP when both pre- and postsynaptic neurons are activated simultaneously while no change in excitatory neurotransmission is evident when only the presynaptic neurons is activated. (G) Computational modeling for the frequency dependent effects of rTMS *in silico*. (H, I) Sample traces and group data of AMPA receptor-mediated miniature excitatory post synaptic currents (mEPSCs) recorded from mouse CA1 pyramidal neurons in control (sham) and stimulated (rMS) cultures 2 – 4 h post stimulation at 1, 5 and 10 Hz (n_sham_ = 14 neurons, n_1Hz_ = 17 neurons, n_5Hz_ = 15 neurons, n_10Hz_ = 14 neurons; Kruskal-Wallis test). Data are mean ± SEM. NS, Not significant. *p<0.05; **p□<□0.01.

### A spike-timing dependent plasticity rule replicates experimental findings *in silico*

Building upon these experimental findings, we employed a computational approach to further explore the mechanisms underlying 10 Hz rMS-induced synaptic plasticity (Figure 3). In these simulations, we employed a previously published STDP rule (14). When examining scenarios in which only afferent inputs were activated, our simulations confirmed the occurrence of LTD in excitatory neurotransmission and successfully replicated LTP induction with simultaneous activation of pre- and postsynaptic neurons (Figure 3E, F); consistent with our observation using optogenetic stimulation. This approach was applied to model rMS-induced plasticity protocols with 900 pulses at 1 Hz; 5 Hz and 10 Hz. The simulations predicted robust potentiation of excitatory neurotransmission at frequencies at 5 Hz and 10 Hz (Figure 3G). To confirm these predictions, experiments were conducted in a separate set of tissue cultures, assessing the effects of stimulation frequency. As shown in Figure 3H and 3I, a significant increase in mean mEPSC amplitude was observed 2 – 4 h with 5 Hz and 10 Hz stimulation.

### 10 Hz rMS induced synaptic plasticity depends on TrkB activation/BDNF signaling

Electrical activation of both pre- and postsynaptic neurons at comparatively low frequencies leads to the release of BDNF from dendrites, a process crucial for inducing STDP (15–18). BDNF plays an important role in synaptic plasticity and several studies have indicated a link between rTMS-induced plasticity and BDNF signaling (19–22). However, no direct experimental evidence exists at the cellular level for a role of BDNF in rTMS-induced synaptic plasticity.

Tissue cultures were stimulated in the presence of a soluble tyrosine receptor kinase B (TrkB/Fc) fusion protein, aimed at scavenging released endogenous BDNF. Indeed, when BDNF was neutralized by the soluble receptor, the 10 Hz rMS-induced increase in mEPSC amplitudes of CA1 neurons was not observed. The absence of an effect from the soluble TrkB/Fc receptor on baseline mEPSC amplitudes supported the idea that endogenous BDNF is released in response to 10 Hz rMS (Figure 4A). Similar results were obtained, when the TrkB receptor was pharamacologically blocked with the TrkB antagonist K252a (100nM).

**Figure 4.**
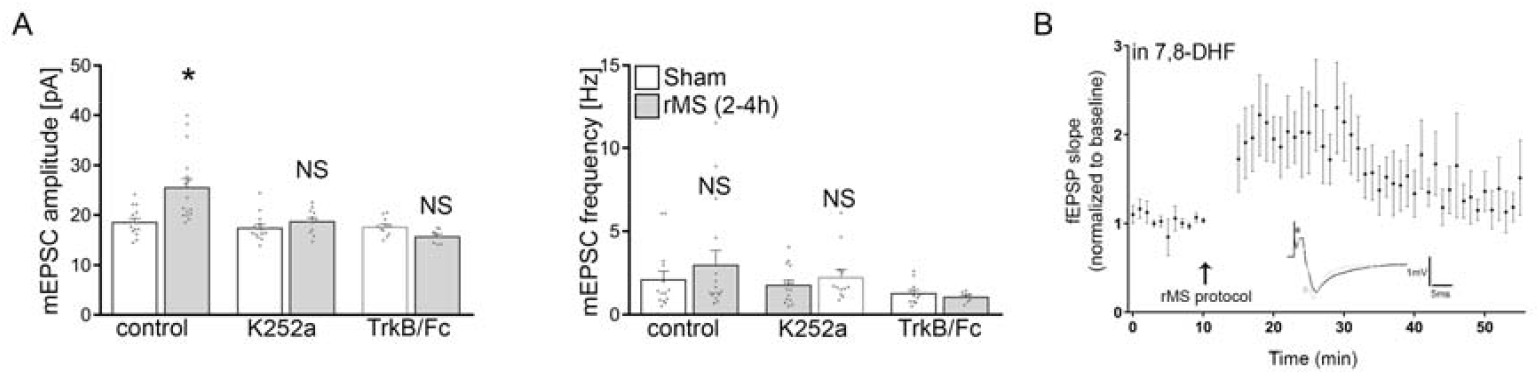
10 Hz rMS induced synaptic plasticity depends on TrkB activation/BDNF signaling (A) Group data of AMPA receptor-mediated miniature excitatory post synaptic currents (mEPSCs) recorded from mouse CA1 pyramidal neurons in control (sham) and stimulated (rMS) cultures 2 – 4 h post stimulation in untreated cultures, cultures that the TrkB receptor was blocked (K252a) and cultures where the BDNF was scavenged using the soluble TrkB/Fc (sham n_control_ = 14 neurons, n_K252a_ = 15 neurons, n_TrkB/Fc_ = 11 neurons; rMS (2-4h) n_control_ = 16 neurons, n_K252a_ = 12 neurons, n_TrkB/Fc_ = 8 neurons; Kruskal-Wallis test). (B) Excitatory post synaptic field potential (fEPSP) slopes recorded in the CA1 stratum radiatum following a 10 Hz, 900 pulses electrical stimulation of the Schaffer collaterals in the presence of the TrkB agonist 7,8-Dihydroxyflavone (7,8-DHF) (n = 4 tissue cultures). Data are mean ± SEM. NS, Not significant. *p⍰<⍰0.05.

Finally, we predicted that activating the TrkB receptor may convert LTD to LTP in experiments where only presynaptic neurons are activated (c.f., Figure 1*G*). To test this hypothesis, we used 10 Hz local electrical stimulation of Schaffer collaterals in the presence of 7,8-dihydroxyflavone (7,8-DHF), a flavone derivative that binds to and activates the TrkB receptor signaling (23, 24). Indeed, the presence of 7,8-DHF during 10 Hz local electrical stimulation resulted in LTP of Schaffer collateral-CA1 synapses (Figure 4B). The results of these experiments suggest that the activation of BDNF/TrkB signaling pathway is essential for rMS-induced plasticity.

## Discussion

The results of this study offer mechanistic insights into the effects of rTMS, demonstrating that coordinated pre- and postsynaptic activation drives synaptic plasticity induced by rMS. We show that an STDP-plasticity rule can predict the frequency-dependent outcome of rTMS. Furthermore, we provide experimental evidence supporting the involvement of TrkB/BDNF signaling in rMS-induced synaptic plasticity. These findings have important implications for understanding the biological effects of rTMS and for developing computational models to predict and standardize the “biological dose” of rTMS.

Over the last decade, rTMS has been increasingly used in both research and clinical practice (25–28). However, the mechanisms underlying rTMS-induced plasticity remain poorly understood. As a result, rTMS protocol design often relies on heuristics, typically based on indirect systems-level measurements or experiments in animal models using (local) electric stimulation. Changes in muscle-evoked potentials during motor cortex stimulation with rTMS in healthy volunteers demonstrated LTP-like plasticity at frequencies ≥ 5 Hz (29–32). In the present study, we confirmed the potentiation of excitatory synapses in organotypic slice cultures at stimulation frequencies ≥ 5 Hz. Our systematic approach utilizing pharmacogenetics, Ca^2+^ imaging, and computational simulations, highlights coordinated pre- and postsynaptic activation as a key mechanism underlying this LTP-like plasticity.

Interestingly, both our simulations and experimental findings revealed no significant changes in excitatory synaptic strength following 1 Hz stimulation. While 1 Hz is known to reduce cortical excitability in humans (38, 39), it has been hypothesized that this frequency might trigger LTD-like plasticity, in line with animal studies using 1 Hz electrical stimulation with 900 pulses to induce LTD in excitatory neurotransmission (33–35). However, the role of STDP has not been fully considered in this context. Our data indicate that 1 Hz rMS does not significantly alter excitatory synaptic strength, suggesting that the inhibitory effects of low-frequency rTMS protocols may not depend on modifications to excitatory synapses. Further investigation is needed to elucidate these processes, while the mechanisms driving rTMS-induced LTP are becoming increasingly well-defined.

BDNF plays a crucial role in synaptic plasticity, including STDP. As a neurotrophin, BDNF regulates synaptic function and plasticity across various neural circuits (36–39), and its activity-dependent release, particularly from dendrites, is essential for plasticity induction (15, 40, 41). Notably, experimental evidence shows that relatively low stimulation frequencies are sufficient to trigger BDNF release when both pre- and postsynaptic compartments are depolarized in a coordinated manner (42). This release is mediated by calcium influx into neurons, which can be triggered by action potentials and synaptic activity. Our previous research has demonstrated that the induction of 10 Hz rMS-LTP requires both NMDA receptor-mediated synaptic calcium influx and L-Type voltage-gated calcium channel-mediated postsynaptic depolarization. Pharmacological blockade of NMDA receptors with APV or L-Type voltage-gated calcium channels with nifedipine prevented the rTMS-induced potentiation of excitatory synapses (4, 9). Interestingly, BDNF adaptively influences synaptic strength through autocrine or paracrine signaling, acting on both the releasing neuron and neighboring cells. Clinically, this suggests a compelling hypothesis: while the exact mechanisms through which rTMS-induced LTP yields positive clinical outcomes remain unclear, it is plausible that rTMS triggers the local release of endogenous BDNF, priming neural network for enhanced synaptic plasticity in response to complex physiological stimuli. This mechanism may also support regeneration following injury and exert neuroprotective effects. Thus, LTP induction may serve as a marker of successful BDNF release, potentially underpinning the therapeutic effects of rTMS, rather than being the direct cause of these effects. Whether BDNF is released during TMS protocols that do not induce LTP, remains an open question and warrants further investigation. Unfortunately, we were unable to reliably detect secreted BDNF in our cultivation medium but application of 7,8-DHF transformed the LTD after presynaptic stimulation only to LTP, suggesting that TrKB receptor activation is crucial for rTMS-induced plasticity. Although 7,8-DHF can also act through other pathways, i.e. PI3K/Akt, ERK, NF-κB and MAPK (43, 44), our control experiments of neutralizing TrkB receptors and scavenging BDNF lead to the same observations. Given recent findings showing changes in serum BDNF levels following TMS stimulation (45–48), we propose that the effective release of BDNF could serve as an important benchmark mechanism for optimizing rTMS protocols. Regardless of these considerations, our experiments provide direct evidence for the role of BDNF in rMS-induced synaptic plasticity. Neutralizing endogenous BDNF with a soluble TrkB receptor blocked 10 Hz rMS-induced LTP, while TrkB receptors activation converted electrically induced LTD into LTP. These findings are consistent with rTMS studies in humans that have indicated an involvement of BDNF in rTMS-induced cortical excitability changes (19, 49). A deeper understanding of the specific signaling pathways facilitating BDNF release during rTMS, combined with a detailed evaluation of stimulation parameters that optimize its release, could open new avenues for therapeutic interventions in neuropsychiatric disorders.

## Materials and Methods

### Ethics statement

Mice were housed under a 12-hour light/dark cycle with unrestricted access to food and water. All efforts were made to minimize animal distress and pain. Experimental procedures adhered to German animal welfare legislation and received approval from the appropriate animal welfare committee and the animal welfare officer at the University of Freiburg.

### Animals

In this study we used C57BL/6J mice of both sexes.

### Preparation of organotypic tissue cultures

Organotypic tissue cultures, 300 μm thick and containing the hippocampus and entorhinal cortex, were prepared from postnatal day 3-5 mice of both sexes as described previously (50). The cultures were maintained in an incubator at 35 °C with 5% CO2 for a minimum of 18 days before experimental assessment. The culture medium was changed three times per week and comprised 50% (v/v) MEM, 25% (v/v) basal medium eagle (BME), 25% (v/v) heat-inactivated normal horse serum, 25 mM HEPES, 0.15% (w/v) NaHCO3, 0.65% (w/v) glucose, 0.1 mg/ml streptomycin, 100 U/ml penicillin, and 2 mM Glutamax, adjusted to pH 7.3 with HCl or NaOH.

### rMS in vitro

Tissue cultures were transferred to a standard 35 mm petri dish containing standard extracellular solution (129 mM NaCl, 4 mM KCl, 1 mM MgCl2, 2 mM CaCl2, 4.2 mM glucose, 10 mM HEPES, 0.1 mg/ml streptomycin, 100 U/ml penicillin, pH 7.4; preheated to 35 °C; 365 mOsm with sucrose). A 70-mm coil (D70 Air Film Coil, Magstim) connected to a Magstim Super Rapid2 Plus1 (Magstim) was positioned 1 mm above the petri dish lid, and the cultures were stimulated using a protocol of 900 pulses at 10 Hz (9 trains of 100 pulses each, 30 s inter-train-interval) at 50 % MSO (maximum stimulator output). The tissue cultures were oriented so that the induced electric field within the tissue was approximately parallel to the dendritic tree of CA1 pyramidal neurons. Time-matched cultures that were not stimulated, but otherwise treated identically, served as controls.

### Whole-cell voltage-clamp recordings

Whole-cell voltage-clamp recordings of CA1 pyramidal neurons were conducted at 35°C as described previously (8). The bath solution consisted of 126 mM NaCl, 2.5 mM KCl, 26 mM NaHCO3, 1.25 mM NaH2PO4, 2 mM CaCl2, 2 mM MgCl2, and 10 mM glucose and was saturated with 95% O2/5% CO2. 10 μM D-APV and 0.5 μM TTX were added in the bath solution to isolate miniature α-amino-3-hydroxy-5-methyl-4-isoxazolepropionic acid receptor-mediated excitatory postsynaptic currents (mEPSCs). The patch pipettes contained 126 mM K-gluconate, 4 mM KCl, 4 mM ATP-Mg, 0.3 mM GTP-Na2, 10 mM PO-creatine, 10 mM HEPES, and 0.1% (w/v) biocytin (pH□7.25 with KOH, 290 mOsm with sucrose). The neurons were recorded at −70□mV. For current-clamp recordings 10 μM D-APV, 10 μM CNQX and 10 μM bicuculline methiodide was added to the bath solution. In some recordings 100 μM clozapine N-oxide (CNO) was washed-in during the recording. For optogenetic stimulation an LED light source was used (pE-300; CoolLED). Series resistance was monitored in 2 – 4□min intervals and recordings were discarded if the series resistance reached ≥30 MΩ and the leak current changed significantly.

### Extracellular field recordings

For extracellular field potential recordings, tissue cultures were placed in a standard interface chamber maintaining a humidified carbogenated (95% O2/5% CO2) atmosphere at 33 ± 1 °C and perfused continuously with artificial cerebrospinal fluid (aCSF) containing 126 mM NaCl, 2.5 mM KCl, 26 mM NaHCO3, 1.25 mM NaH2PO4, 2 mM CaCl2, 2 mM MgCl2, and 10 mM glucose and saturated with 95% O2/5% CO2. A glass recording electrode containing 0.75M NaCl was placed in the *stratum radiatum* of CA1 while a bipolar electrode was placed approximately 1 mm proximal to the recording pipette to stimulate the Schaffer collaterals. Baseline and post-stimulation values were recorded at a frequency of 0.033 Hz. In some experiments, stimulation of virus expressing cells was done using an LED light source (pE-300; CoolLED).

### Viral transduction

Tissue cultures were transfected between 3 and 6 days in vitro (DIV) by adding 1□μl of AAV-hSyn1-GCaMP6f-P2A-nls-dTomato [gift from Jonathan Ting (Addgene viral prep # 51085-AAV1; http://n2t.net/addgene:51085 ; RRID:Addgene_51085)] directly on top of each culture. For local viral transduction approximately 100 nL of AAV-hSyn-hM4D(Gi)-mCherry [gift from Bryan Roth (Addgene viral prep # 50475-AAV9 ; http://n2t.net/addgene:50475 ; RRID:Addgene_50475)] or AAV-Syn-ChrimsonR-tdT [gift from Edward Boyden (Addgene viral prep # 59171-AAV9; http://n2t.net/addgene:59171 ; RRID:Addgene_59171] were injected within the area of interest using a glass pipette with the help of a micromanipulator. Tissue cultures were then left to mature at least until 18 DIV before experimental assessment.

### Calcium imaging

Filter inserts containing transduced slices were transferred to a custom-built carbon bridge at an upright fluorescent microscope (LN Scope; Luigs and Neumann) equipped with a 10x water immersion objective (NA 0.3; Olympus). The image acquisition was performed using an ORCA-Fusion camera (Hamamatsu Photonics) and a Lambda DG-4 Plus as a light source (Sutter Instruments). The bath solution contained NaCl 129 mM, KCl 4 mM, MgCl2 1 mM, CaCl2 2 mM, glucose 4.2 mM, HEPES 10 mM, Trolox 0.1 mM, streptomycin 0.1 mg/ml, penicillin 100 U/ml (pH 7.4; at 35°C). The D70 AirFilm Coil (Magstim) was placed directly underneath the recording chamber oriented in a way that the induced electric field was parallel to the dendritic trees of CA1 pyramidal neurons. The stimulation protocol consisted of 900 pulses at 10 Hz at 50% MSO. Image acquisition and data analysis was performed with the MetaFluor® software (Molecular Devices).

### *Post hoc* identification of recorded neurons and imaging

Tissue cultures were fixed in a solution of 4% (w/v) PFA and 4% (w/v) sucrose in 0.01 M PBS for 1 h. Fixed cultures were incubated for 1 h with 10% (v/v) NGS and 0.5% (v/v) Triton X-100 in 0.01 M PBS. Biocytin filled cells were stained with appropriate Alexa-conjugated streptavidin (Thermo Fisher Scientific; 1:1000; in 0.01 M PBS with 10% NGS and 0.1% Triton X-100) for 4 h, and TOPRO or DAPI (Thermo Fisher Scientific) staining was used to visualize cytoarchitecture (1:5000; in 0.01 M PBS for 15□min). Tissue cultures that were transduced were only stained with DAPI after fixation. Slices were washed, transferred, and mounted onto glass slides for visualization. Transfected and streptavidin-stained neurons were visualized with a Leica Microsystems TCS SP8 laser scanning microscope with 20× (NA 0.75), 40× (NA 1.30), and 63× (NA 1.40) oil-submersion objectives.

### Computational modeling

For the simulation of rTMS effects on neurons a simple model was used (14). This model consists of two identical one-compartment neurons connected by an excitatory synapse. The neuronal dynamics are a modified version of the Hodgkin-Huxley model and represent a minimal form of regular-spiking (RS) neurons (51). The synapse is modeled as an excitatory alpha synapse, incorporating both short-term plasticity (STP) and spike-timing-dependent plasticity (STDP). The model was optimized by the addition of random spikes with an average frequency of 1 Hz for both neurons, a synaptic weight of 0.01 μS, a synaptic delay of 0.5 ms, and STDP parameters were set as follows: hebbw: 0.00002, antiwt: -0.00002, Tauhebb: 80, and Tauanti: 80. Source parameters included a type of IClamp, with an amplitude of 137% of the threshold for the pre-synapse and 125% of the threshold for the post-synapse. All other parameters remained as were (14). Using these settings, 900 pulses at 10 Hz of electric current were applied through the I-clamp to mimic rTMS pulses on either the pre- or post-synaptic neurons. The maximum conductance of the synapse was measured during stimulation, while data were recorded for 60 s before and 180 s after stimulation. All simulations were performed using NetPyNE version 1.0.1 (52), and the code for all simulations is available at https://github.com/mrvasheghani/rMS-induces-cooperative-plasticity.

### Experimental design and statistical analysis

Analyses were performed with the person analyzing the data blind to the experimental condition. For this project, we used one or two tissue cultures from each animal. Electrophysiological data were analyzed using pClamp 11.2 software suite (Molecular Devices) and Easy Electrophysiology 2.5.0.2 (Easy Electrophysiology Ltd.). Statistical comparisons were made using Mann–Whitney test (to compare two groups) two-way ANOVA and Kruskal-Wallis test as indicated in the figure captions and text (GraphPad Prism 7). p values of <0.05 were considered a significant difference. All values represent mean ± SEM.

### Digital illustrations

Confocal image stacks were exported as 2D projections and stored as TIFF files. Figures were prepared using Photoshop graphics software.

## Acknowledgments

The authors thank Rodrigo Alejandro Perez for support in the field potential recordings, and Emina Deumic and Susana Glaser for their skillful assistance in tissue culturing. This work was supported by the Federal Ministry of Education and Research, Germany (BMBF, 01GQ2205A).

